# The Immunological Proteome Resource

**DOI:** 10.1101/2022.08.29.505666

**Authors:** Alejandro J. Brenes, Jens L. Hukelmann, Laura Spinelli, Andrew J.M. Howden, Julia M. Marchingo, Linda V. Sinclair, Christina Rollings, Olivia J. James, Iain R Phair, Stephen P. Matthews, Sarah H. Ross, J. Simon C. Arthur, Mahima Swamy, David K. Finlay, Angus I. Lamond, Doreen A. Cantrell

## Abstract

The Immunological Proteome Resource (ImmPRes; http://immpres.co.uk/) is an open access public resource integrating proteomic data generated by large-scale mass-spectrometry analysis of murine hematopoietic populations. The initial focus is T lymphocytes and how their proteomes are shaped by immune activation, environment, and intracellular signalling pathways with an aim to expand it to B cells and innate immune cells. It is a multidisciplinary effort between immunology and mass spectrometry-based labs with the objective to help define an in-depth high-quality map of immune cell proteomes. Maintaining data reproducibility and integrity are a priority within the resource, thus there is an in-depth protocols section explaining in detail the sample processing and the mass spectrometry-based analysis. ImmPRes provides open access to proteomic datasets covering a wide range of murine leukocyte populations with analysis of copy numbers per cell of > 10,000 proteins, enabling new understanding of lymphocyte phenotypes. All data is accessible via a simple graphical interface that supports easy interrogation of the data and options to download data summaries and raw data files.

## Background and summary

The immune system is a vast network of interacting cells that function to protect organisms against foreign pathogens. Understanding the roles of different cells, signalling pathways and proteins within this system is vital to understanding health and disease. In this context, high quality RNA based resources^1-3^ have played vital roles in defining expression levels of RNAs across bulk and single cell immune populations. However, proteins are the molecules that form the backbone of cell structure and control virtually all metabolic processes and regulatory mechanisms^4^. The proteome of a cell is a dynamic system that is constantly modulated by changes in rates of protein synthesis and degradation^5^. mRNA levels are thus not effective predictors of protein abundance^6,7^, and quantitative characterisation of cellular proteomes versus transcriptomes has been shown to provide invaluable information about lymphocyte identity^8^.

Mass spectrometry instrumentation, sample processing and software level break-throughs have enabled the comprehensive characterisation of the proteome in a scalable and cost-effective manner^9^. Thus, defining the proteome of hematopoietic cells in steady state, as well as in response to different conditions such as T cell receptor activation or specific cytokines has become an attainable project of great potential benefit to the immunological community. A resource looking at a select number of human hematopoietic populations at the protein level was recently developed^10^, but there is no current resource exploring the proteomes of murine leukocyte populations. Mice are widely accepted to be excellent models to study the mammalian immune system^11^ and have played a vital role in immunology research. The tractability of mouse genomes means that mice have been used extensively to explore the importance of different immune cell populations and different immune regulatory molecules. As such, resources mapping the proteomic profile of murine hematopoietic cells and in particular mouse T cell populations would be of great value to the community.

To achieve this goal, we created The Immunological Proteome Resource (ImmPRes; http://immpres.co.uk/), an open access public resource integrating proteomic data generated by large-scale mass-spectrometry analysis of murine hematopoietic populations. It is a multidisciplinary collaborative effort between immunology and mass spectrometry-based labs with the objective to help define an in-depth high-quality map of the immune proteome.

One of the main goals of ImmPRes is to provide open access to all proteomic data, ranging from the raw MS files to the processed copy number summaries. It does so by providing access to a simple graphical interface designed to interact with the different proteomic datasets, and to download the raw data for large-scale reanalysis. All MS raw files are uploaded to Proteomics Identifications Database^12,13^ (PRIDE) where they can be downloaded and reanalysed, while the processed data is made available for download directly on ImmPRes.

Data reproducibility and integrity are a priority within the resource. As such there is an in-depth protocols section explaining in detail both the sample processing as well as the mass spectrometry analysis. Furthermore, the sample preparation steps for each of the different hematopoietic populations are documented within the protocols section. Specific details on the mouse strains, tissue preparation protocols, growth media, cytokines added, purification protocols and activation details (where applicable) are available within the protocols section. The current mass spectrometry-based datasets have been acquired in Data Dependent Acquisition (DDA) mode using either an SP3^14^ or urea based sample preparation coupled with isobaric labelling using Tandem Mass Tags^15^ (TMT) (Fig. 1a), or a label-free strategy (Fig. 1b). All the DDA data were acquired with extensive fractionating of the samples for an in-depth overview and were analysed with a rigorous false discovery rate and with data imputation disabled. Furthermore, all datasets containing heterogenous populations were searched without using “match between runs” to avoid false identifications being propagated^16^. For the analytic side, the “proteomic ruler”^17^ method was used for the normalisation of all current datasets. This method uses the mass spectrometry signal of histones as an internal standard which avoids the error prone steps of cell counting and enables the estimation of protein copy numbers per cell. Knowledge of protein copy number is invaluable for a full understanding of cell phenotypes. For example, it allows exploration of the stoichiometry of proteins within protein complexes and provides an abundance measure with direct biological meaning to the end user.

**Figure 1.**
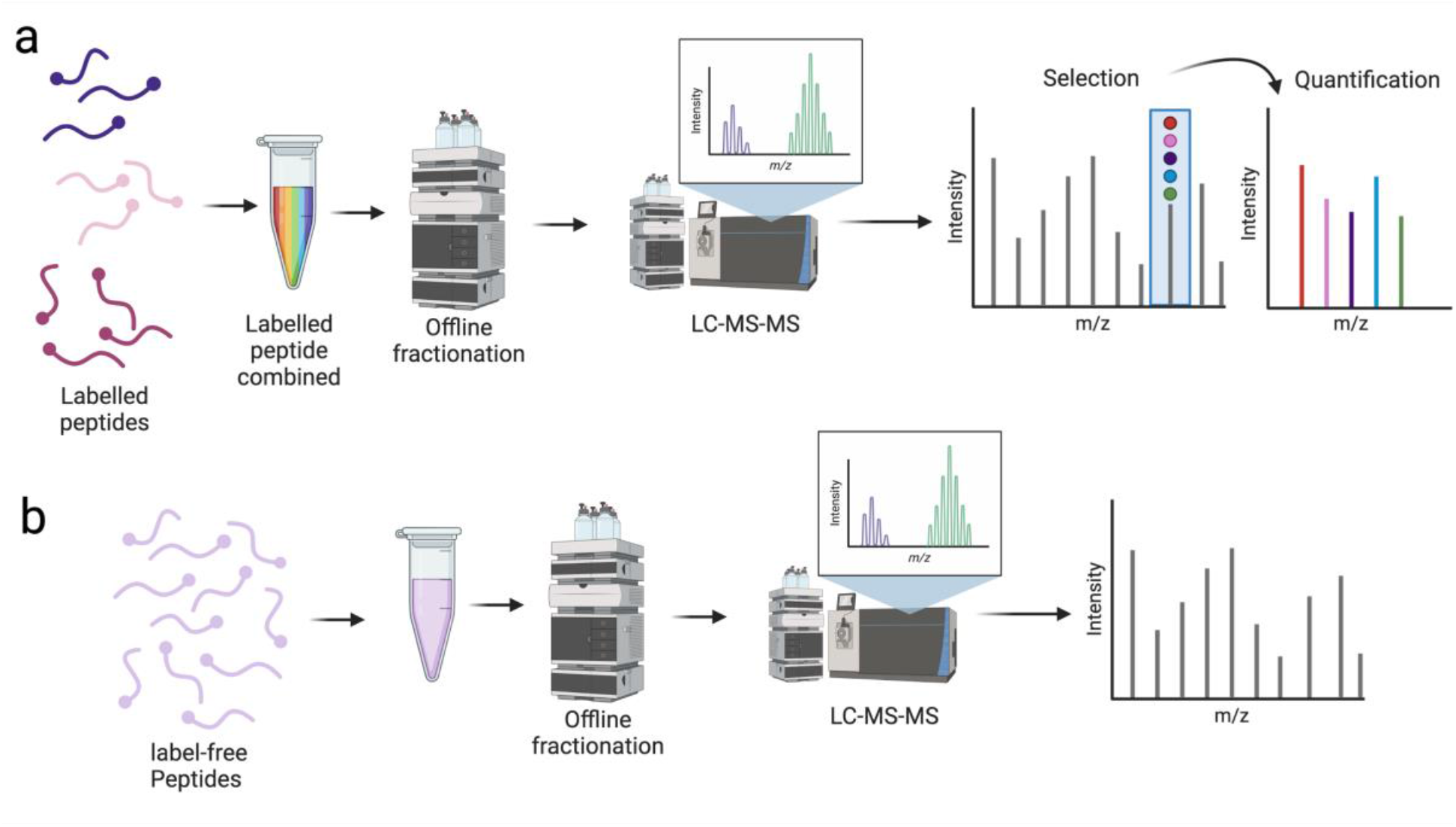
Mass spectrometry workflows: ***(a)*** Schematic showing the SPS-MS3 TMT workflow that was used for all TMT datasets. ***(b)*** Schematic showing the label free workflow that was used for the label free datasets.

## Data Records

ImmPRes contains multiple large-scale proteomics datasets containing hundreds of raw files and identifying thousands of proteins. The initial resource has integrated numerous datasets (Fig. 2), across different hematopoietic populations and across different conditions. The data is readily available to be browsed and explored using the graphical interface.

**Figure 2.**
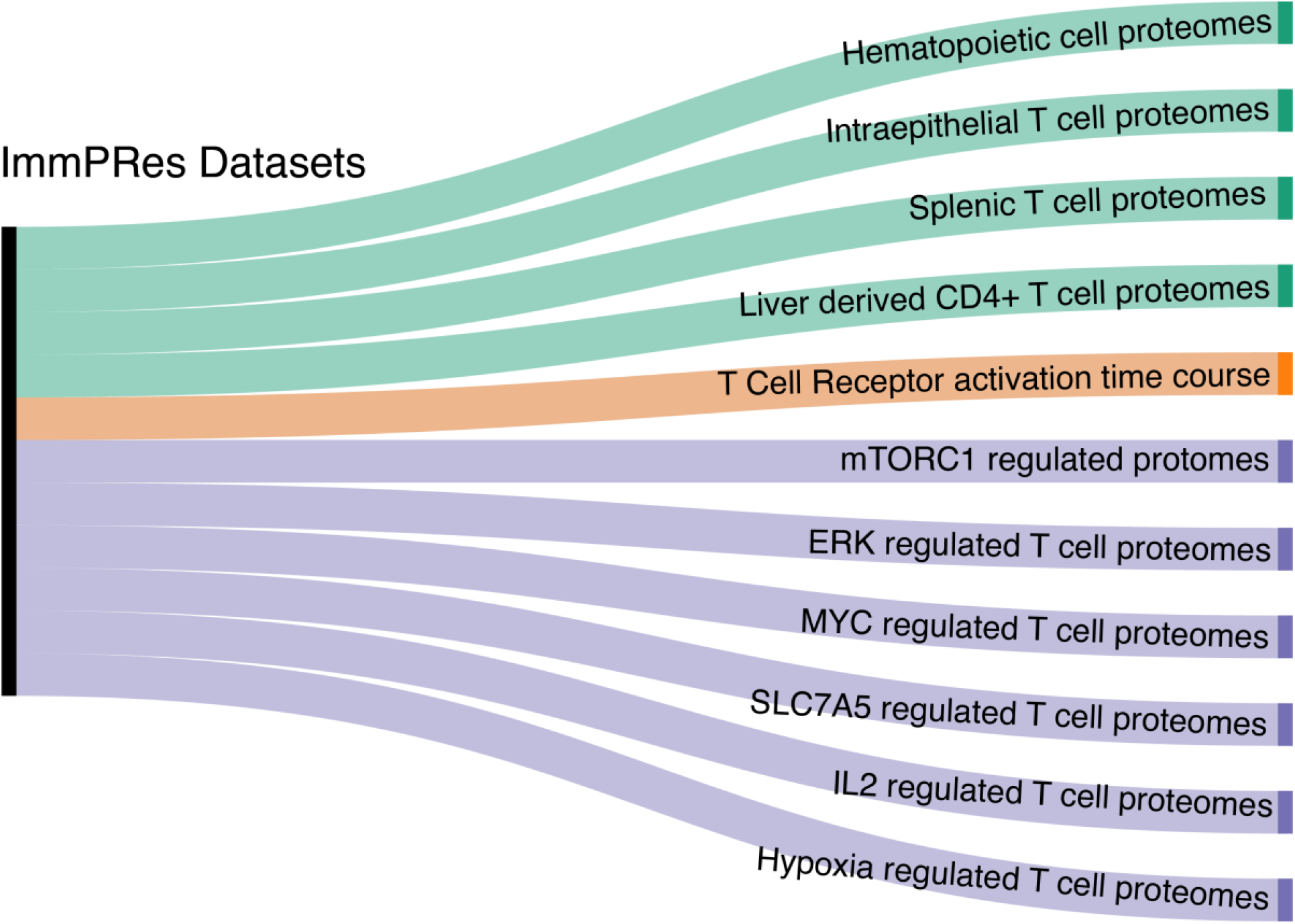
Overview of all data records

### Hematopoietic cell proteomes

The dataset labelled **‘Hematopoietic cell proteomes’** is the largest and most in-depth dataset hosted on ImmPRes. It is a TMT-based liquid chromatography-mass spectrometry (LC-MS) characterization of 16 different immune cell populations, including 8 previously unpublished populations (Fig. 3a). This large dataset contains naïve CD4+ and CD8+ T cell subpopulations isolated from lymph nodes, multiple in vitro generated effector CD4 T cell subsets, effector and memory like CD8 subpopulations, in vitro generated CD4+ regulatory T cells well as innate leukocytes such as mast cells and macrophages^18^. The in-depth coverage meant that over 10,000 proteins were identified across the whole dataset, without using matching or imputation (Fig. 3b). Furthermore, all populations had a peptide coverage greater than 55,000 peptides and ranging up to almost 99,000 peptides (Fig. 3c). The raw files and MaxQuant output for this dataset are available in PRIDE with identifier PXD012058^19^ and PXD020091^20^ while the processed copy numbers are available in the ‘Downloads’ tab in ImmPRes.

**Figure 3.**
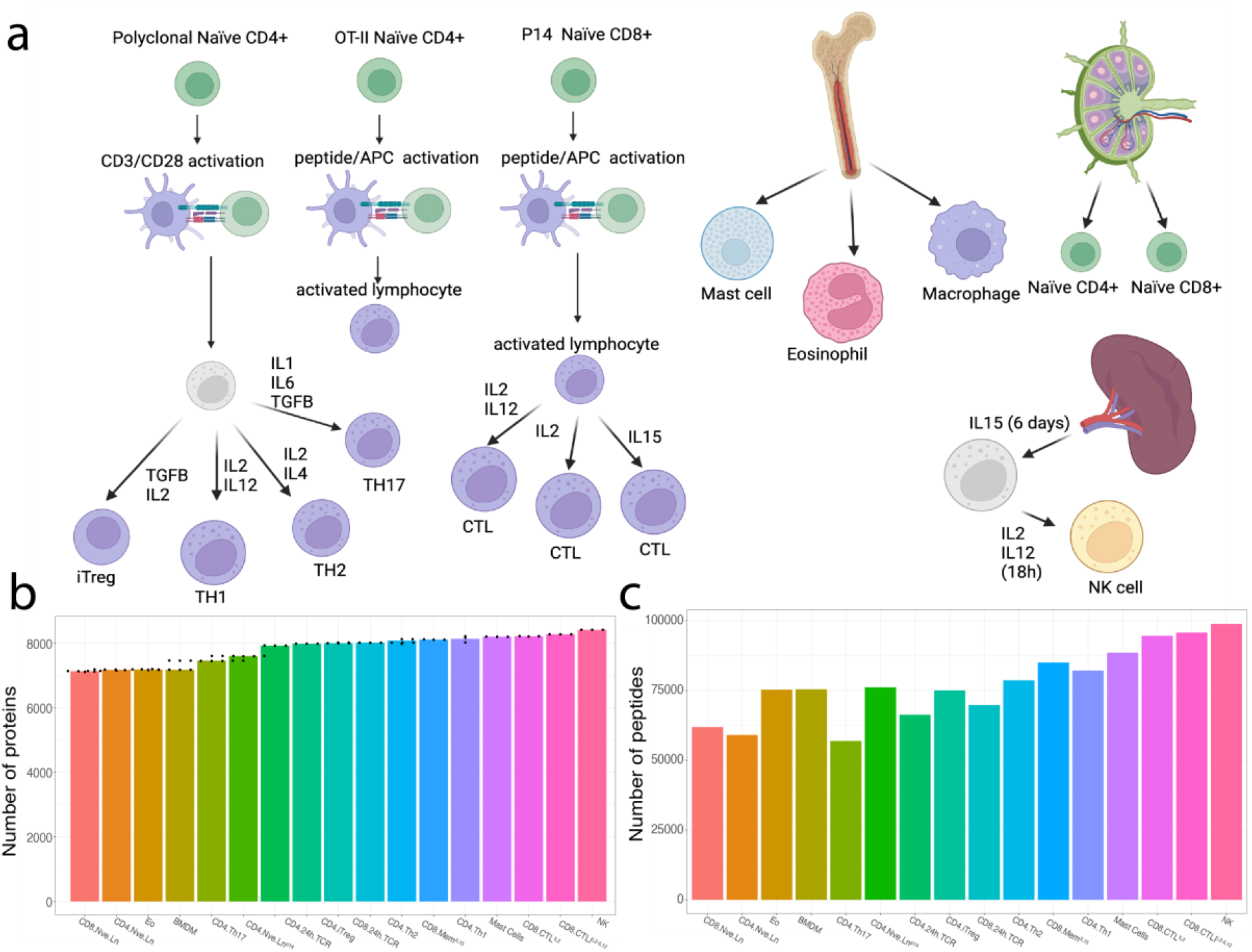
Primary data record: ***(a)*** Schematic showing the different populations which are contained within the ‘Hematopoietic cell proteomes’ data record. ***(b)*** Bar plot showing the number of proteins identified for all populations. ***(c)*** Bar plot showing the number of peptides identified for all populations.

### Splenic T cell proteomes

T cells in the spleen are frequently used to study T cell biology. ImmPRes includes a TMT-based LC-MS dataset, labelled as **‘Splenic T cell proteomes’** on the Data Browser, characterising the proteomes of ex-vivo splenic T cells, including naïve CD8+ T cells, naïve CD4+ T cells, memory like CD4+ T cells and CD4+ regulatory T cells (Fig. 4a). The raw files and MaxQuant output for this dataset are available in PRIDE with identifier PXD020091^20^, while the processed copy numbers are available in the ‘Downloads’ tab in ImmPRes.

**Figure 4.**
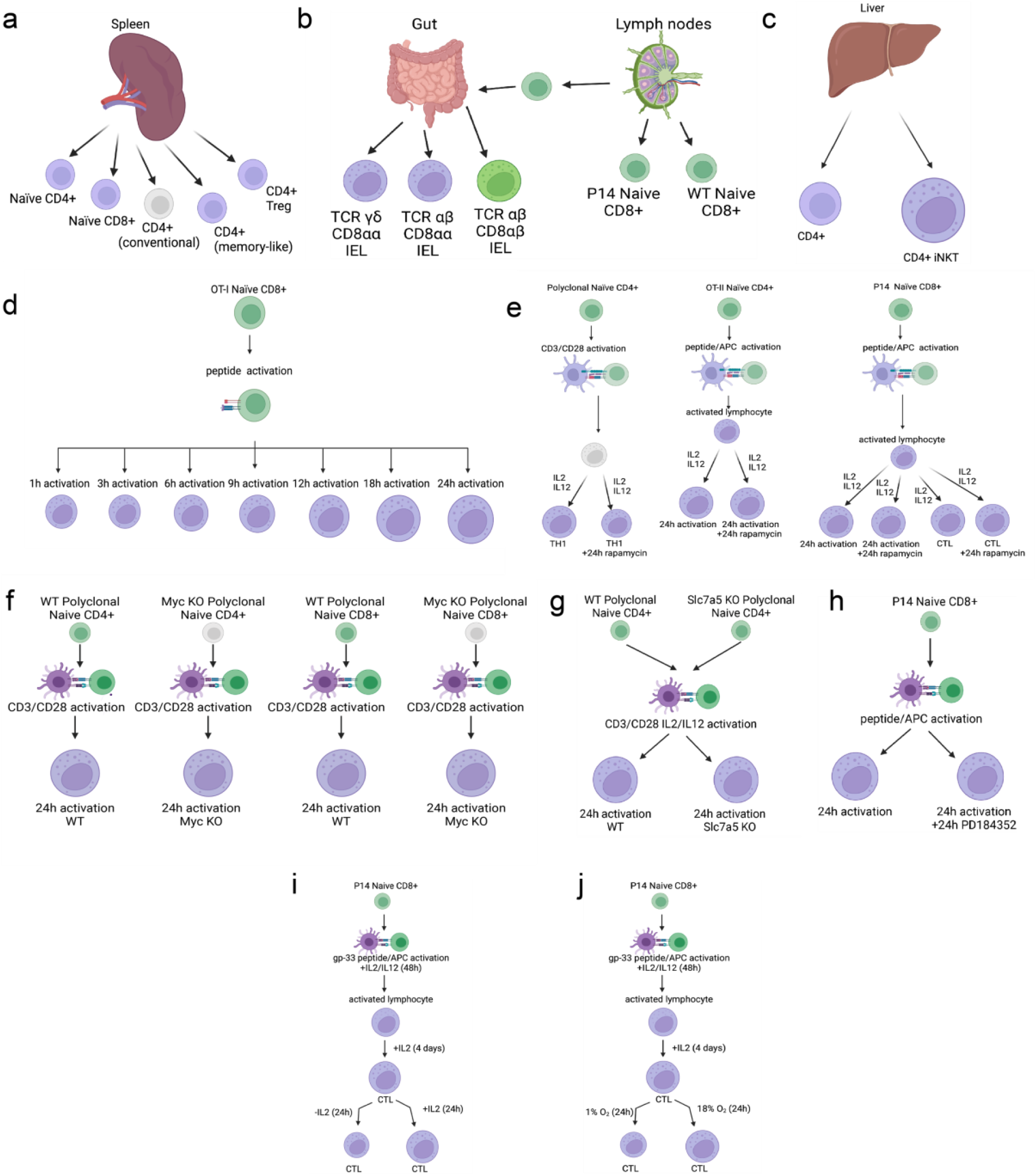
Data records overview: Schematics representing: ***(a)*** Populations contained in the ‘Splenic T cell proteomes’ data record. ***(b)*** Populations which are contained within the ‘Intraepithelial T lymphocyte proteomes.’ data record. ***(c)*** Populations contained within the ‘Liver derived CD4+ T cell proteomes.’ data record. ***(d)*** Populations contained within the ‘T Cell Receptor activation time course.’ data record. ***(e)*** Populations and treatments contained within the ‘Mtorc1 regulated proteome’ data record. ***(f)*** Populations contained within the ‘Myc regulated proteomes’ data record. ***(g)*** Populations contained within the ‘Slc7a5 regulated proteomes’ data record. ***(h)*** Populations and treatments contained within the ‘Erk regulated proteomes’ data record. ***(i)*** Populations and treatments contained within the ‘IL2 regulated proteomes’ data record. ***(j)*** Populations and treatments contained within the ‘Hypoxia regulated proteomes’ data record.

### Intraepithelial T lymphocyte proteomes

Intestinal intraepithelial lymphocytes (IEL) comprise a distinct group of innate-like and memory T cells that collectively form one of the largest T cell compartments in the body. ImmPRes includes a TMT-based LC-MS dataset exploring the proteomes of intraepithelial T lymphocytes (T-IELs), labelled as **‘Intraepithelial T cell proteomes’** on the Data Browser (Fig. 4b). The dataset characterises the proteome of TCRαβ CD8αβ, TCRαβ CD8αα and TCRγδ CD8αα T-IELS along with two conventional TCRαβ CD8αβ lymph node derived naïve CD8+ T cell populations, wild type (WT) and P14^21^. The raw files and MaxQuant output for this dataset are available in PRIDE with identifier PXD023140^22^, while the processed copy numbers are available in the ‘Downloads’ tab in ImmPRes.

### Liver derived CD4+ T cell proteomes

T cells play a critical role in liver immunity and take part both in the initiation and in the resolution of intrahepatic inflammation^23^. The liver contains conventional CD4 T cells, and Natural Killer T (NKT) cells that express an invariant Vα14 T cell receptor that recognizes glycolipid/CD1d antigen complexes (iNKTs) and play a role in immune surveillance and immune homeostasis^23^. ImmPRes includes a TMT-based LC-MS dataset, labelled as **‘Liver derived CD4+ T cell proteomes’** on the Data Browser, characterising the proteomes of ex-vivo liver derived CD4+ T cells along with iNKT cells (Fig. 4c). The raw files and MaxQuant output for this dataset are available in PRIDE with identifier PXD036319^24^, while the processed copy numbers are available in the ‘Downloads’ tab in ImmPRes.

### Time course analysis of CD8+ T cell proteome remodelling in response to T Cell Receptor (TCR) engagement

A label-free LC-MS dataset, labelled as **‘T Cell Receptor activation timecourse’** on the Data Browser, analysing the proteome of CD8+ naïve T cells responding to cognate antigen over a time course (Fig. 4d). The T cells were CD8+ cells expressing a TCR complex that recognises the ovalbumin peptide SIINFEKL in the context of H2Kb (OT1-T cells). The time course data was collected at 0, 1, 3, 6, 9, 12, 18 and 24 hours of activation of OT-1 T cells with SIINFEKL. The raw files and MaxQuant output for this dataset are available in PRIDE with identifier PXD016443^25^, while the processed copy numbers are available in the ‘Downloads’ tab in ImmPRes.

### mTORC1 regulated lymphocyte proteomes

One key signalling molecule that controls protein turnover in mammalian cells is the nutrient sensing protein kinase mammalian target of rapamycin complex 1 (mTORC1)^26^. ImmPRes contains a TMT-based LC-MS dataset, labelled as ‘**mTORC1 regulated proteomes**’ on the Data Browser, which explores the effect of how rapamycin, an inhibitor of mTORC1 reshapes T cell proteomes (Fig. 4e). The data compares how mTORC1 inhibition impacts the immediate response of naïve CD4+ and CD8+ T cells to antigen, versus how 24 hours of mTORC1 inhibition reshapes differentiated effector CD4+ TH1 cells and CD8+ T cytotoxic T lymphocytes (CTL)^18^. The impact of mTORC1 inhibition on six additional populations are set to be released in the near future. The raw files and MaxQuant output for this dataset are available in PRIDE with identifier PXD012058^19^ and PXD020091^20^, while the processed copy numbers are available in the ‘Downloads’ tab in ImmPRes.

### Myc control of the proteomic landscape of immune activated CD4+ and CD8+ T cells

T cell expansion and differentiation are critically dependent on the transcription factor Myc^27^. ImmPRes contains a label-free LC-MS dataset, labelled as ‘**Myc regulated T cell proteomes**’ on the Data Browser, studying the effects of Myc-deletion on the proteomes of immune activated CD4+ and CD8+ T cells (Fig. 4f). The dataset comprises wild type CD4+ and CD8+ naïve T cells, along with wild type and Myc null CD4+ and CD8+ T cells polyclonally activated with CD3 and CD28 antibodies^27^. The raw files and MaxQuant output for this dataset are available in PRIDE with identifier PXD016105^28^, while the processed copy numbers are available in the ‘Downloads’ tab in ImmPRes.

### Analysis of how the amino acid transporter SLC7A5 fuels CD4+ T cell proteomes

T lymphocytes regulate nutrient uptake to meet the metabolic demands of an immune response. They respond to antigen by upregulating expression of many amino-acid transporters, including SLC7A5, the System L (‘leucine-preferring system’) transporter. SLC7A5 transports large neutral amino acids into T cells and is the dominant methionine transporter^29^, accordingly SLC7A5 null T cells cannot proliferate or differentiate in response to antigen. ImmPRes includes a label-free LC-MS dataset, labelled as **‘SLC7A5 regulated T cell proteomes’** on the Data Browser, showing the effects of Slc7a5-deficiency on the proteomes of immune activated CD4+ T cells (Fig. 4g). The dataset compares WT naïve CD4+ T cells and wild type and SLC7A5 null CD4+ T cells polyclonally activated with CD3 and CD28 antibodies in the presence of cytokines interleukin 2 (IL2) and IL12^27^. The raw files and MaxQuant output for this dataset are available in PRIDE with identifier PXD016105^28^, while the processed copy numbers are available in the ‘Downloads’ tab in ImmPRes.

### Analysis of how antigen receptor driven proteome restructuring in CD8+ T cells is regulated by extracellular signal-regulated kinases (ERKs)

A key T cell signalling module is mediated by ERK1/2 serine/threonine kinases which are activated in response to antigen receptor engagement. ImmPRes contains a label**-**free LC-MS dataset, labelled as **‘ERK regulated T cell proteomes’** on the Data Browser, showing the effect of inhibiting ERK activity on antigen receptor driven proteome restructuring in CD8+ T cells (Fig. 4h). The dataset compares naïve CD8+ T cells and CD8+ T cells expressing antigen receptors specific for lymphocytic choriomeningitis virus glycoprotein peptide gp33-41 activated for 24 hours with the gp33 peptide in the presence or absence of the kinase inhibitor PD184352 which prevents activation of ERK1/2^30^. The raw files and MaxQuant output for this dataset are available in PRIDE with identifier PXD023256^31^, while the processed copy numbers are available in the ‘Downloads’ tab in ImmPRes.

### Analysis of how Interleukin 2 controls the proteome of CD8 cytotoxic T cells

Interleukin-2 (IL-2) regulates transcriptional programs and protein synthesis to promote the differentiation of effector CD8^+^ cytotoxic T lymphocytes (CTLs)^32^. ImmPRes contains a label**-** free LC-MS dataset, labelled as **‘IL2 regulated T cell proteomes’** on the Data Browser, showing the effect of IL2 on CTL expressing T cell receptor specific for the gp33 peptide from lymphocytic choriomeningitis virus (Fig. 3h). P14 naïve CD8+ T cells were activated with the gp33-41 peptide for 48 hours and cultured in IL2 for 4 days to generate CTL. A population of CTLs were maintained in IL-2, while another was deprived of IL-2 for 24 hours^33^. The raw files and MaxQuant output for this dataset are available in PRIDE with identifier PXD008112^34^, while the processed copy numbers are available in the ‘Downloads’ tab in ImmPRes.

### Analysis of hypoxia induced remodelling of cytotoxic T cell proteomes

During immune responses T cells must function in oxygen-deficient, or hypoxic, environments. ImmPRes contains label-free LC-MS dataset, labelled as **‘Hypoxia regulated T cell proteomes’** on the Data Browser, showing the effect of 24 hours of hypoxia (1% oxygen saturation) on the proteomes of CTL (Fig. 3i). P14 naïve CD8+ T cells were activated with the gp-33 peptide for 48 hours and cultured in IL2 for 4 days to generate CTL. A population of CTLs was then maintained in hypoxic conditions (1% oxygen saturation) for 24 hours, while another was maintained in normoxic conditions (18% oxygen saturation) for 24 hours^35^. The raw files and MaxQuant output for this dataset are available in PRIDE with identifier PXD026223^36^, while the processed copy numbers are available in the ‘Downloads’ tab in ImmPRes.

## Technical validation

All the of technical details relating to the staining, isolation, activation and or treatment procedures for all the different populations are included within the ‘Protocols & publications’ tab. For all samples processed and subject to mass spectrometry, the EZQ assay was used to determine a suitable protein yield. Furthermore, additional quality control (QC) steps included running blanks between the different samples to avoid signal carry over and running a commercial QC standard (HeLa lysates) to ensure the instrument performance was within suitable parameters.

All datasets were processed with MaxQuant^37,38^, but to avoid the issue of false protein identifications induced by lack of FDR control within ‘Match-between-runs’^16^, this setting was disabled for all datasets containing heterogenous populations. Furthermore, there has been no use of data imputation across any population or dataset. On the quantification side, the proteomic ruler^17^ was the selected method for all populations, as such all datasets were checked to contain a peptide depth greater than 12,000 to ensure they would be suitable to the requirements of the method. All proteins which were labelled as ‘Reverse’, ‘Contaminants’ or ‘Only identified by site’ by MaxQuant were filtered out from the results. Proteins identified across different populations by a single peptide are included within the results with a cautionary message regarding their reduced reliability.

## Usage notes

All data hosted on ImmPRes are open access under CC BY 4.0 terms. As a web resource with a user-friendly graphical interface, users can easily generate interactive plots. This functionality is present for all datasets and can be found under the ‘Data browser’ tab. Within the Data browser the first sub tab labelled ‘Datasets’ provides a graphical representation of the different collections of data that have been integrated into ImmPRes, these are also shown in Figures 2-4.

The **‘Copy numbers’** tab (Fig. 5a) is a simple way to explore the estimated copy numbers for a particular protein of interest in a specific dataset. This tab has multiple dropdowns with important functionality, the first dropdown is labelled **‘Dataset selection’** and it allows users to select which datasets they want to visualise from all available options in ImmPRes. This field is particularly informative as it will display the dataset name, i.e. ‘Hematopoietic cell proteomes’, it will also state the acquisition method used for the dataset, DDA (data dependent acquisition) or DIA (data independent acquisition), and it will show the labelling strategy used, be it Tandem Mass Tags or Label-free. The **‘Protein search (gene name based)’** dropdown allows users to search for their protein of interest using the standard Gene name, the default option shows the results for ‘Pten’. Additionally, some datasets allow the user to filter the results by selecting specific populations, i.e. within the ‘Hematopoietic cell proteomes’ dataset users can filter the results to show only ‘CD8+ T cells’.

**Figure 5.**
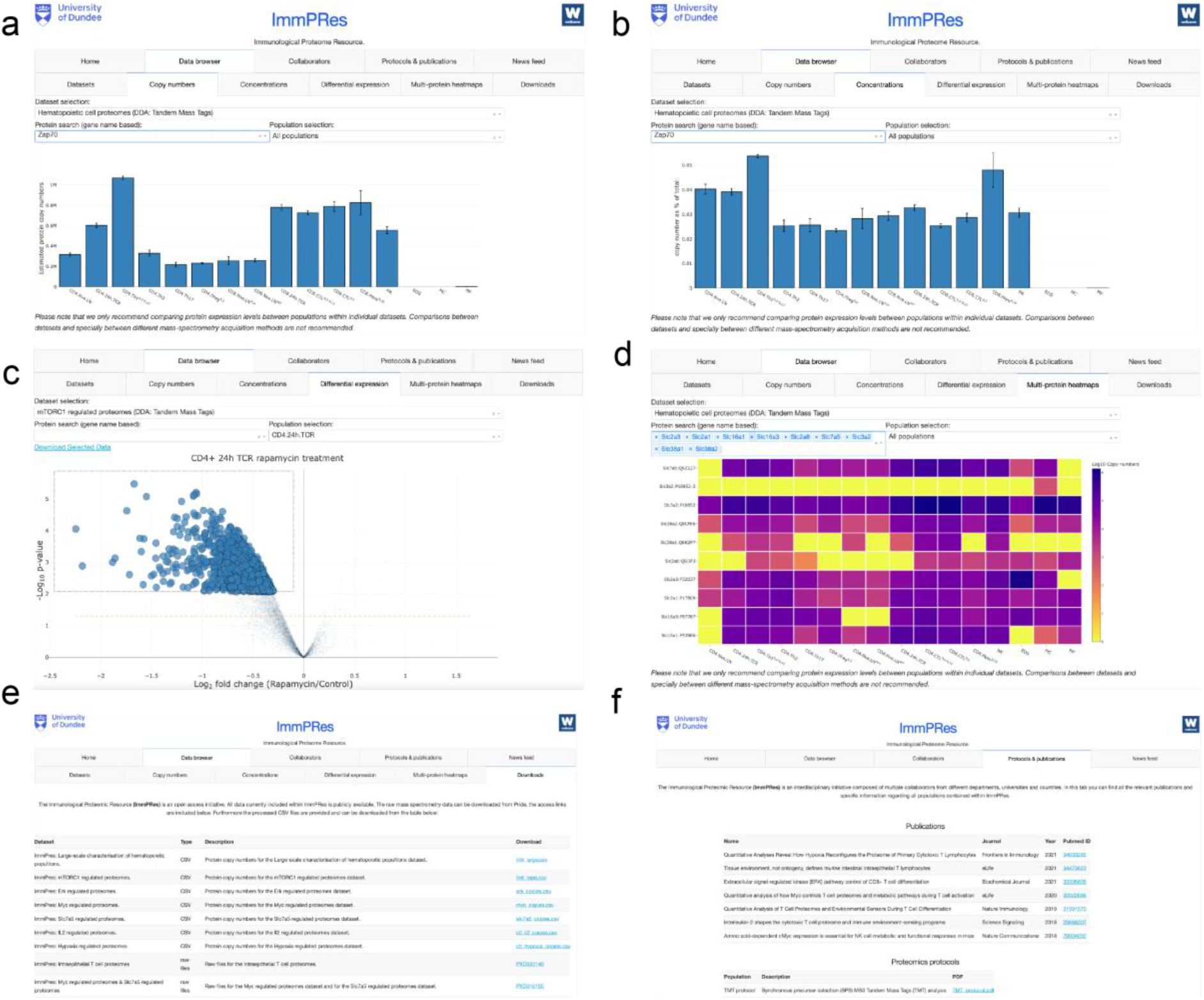
Functionality: **(a)** Snapshot showing the ‘Copy numbers’ tab within immpres.co.uk. Different proteins can be searched by typing their gene name within the ‘Gene Search’ box. **(b)** Snapshot showing the concentrations tab within immpres.co.uk. Different proteins can be searched by typing their gene name within the ‘Gene Search’ box. **(c)** Snapshot showing the ‘Differential expression’ tab within immpres.co.uk. Different proteins can be searched by typing their gene name within the ‘Gene Search’ box. **(d)** Snapshot showing the ‘Multi-protein heatmaps’ tab within immpres.co.uk. Different proteins can be searched by typing their gene name within the ‘Gene Search’ box. **(e)** Snapshot showing the ‘Downloads tab within immpres.co.uk **(f)** Snapshot showing the ‘Protocols & publications’ tab within immpres.co.uk

The **‘Concentrations’** tab (Fig. 5b) mimics the previously described behaviour, but instead of showing the estimated copy numbers, it will show a concentration-like measure (copy numbers of a protein divided by the total copy numbers), providing a measurement that normalises differences in cell size, it is an alternative way to visualise the expression levels of proteins across the datasets. The next tab within the Data Browser is the **‘Differential expression’** tab (Fig. 5c). This tab provides access to a volcano plot; a plot which shows the fold change and p-values when comparing two specific conditions. The volcano plots are available for some datasets and for some specific comparisons. Like the two previous tabs, the Differential expression tab has a **‘Dataset selection’** dropdown, a ‘**Protein search (gene name based)**’ dropdown and a **‘Population selection’** dropdown. Users can download the results shown on the volcano plot by clicking on the ‘Download Selected Data’ button. Users can also select a segment of the volcano by pressing and holding the left click over the desired section in the plot and can download this subset of proteins by pressing the ‘Download Selected Data’ button.

The final interactive tab is the **‘Multi-protein heatmaps’** (Fig. 5d). The functionality of this tab is the same as described above, except for the ‘**Protein search (gene name based)**’ dropdown, which unlike previous tabs allows the users to search for multiple proteins at the same time. Every time a new protein is selected it will update the heatmap to also include the new selection. The heatmap itself shows the log_10_ converted protein copy numbers for the selected proteins and populations within the selected dataset.

The Data browser also provides access to the data should users wish to explore any dataset outside of ImmPRes. Within the ‘Downloads’ tab (Fig. 5e) there is a table that lists all the processed data, with comma separated value (CSV) files to download the estimated copy numbers for all datasets, as well as links to PRIDE^12^ which allow users to download the raw MS files for reanalysis. The final tab of relevance to usage is the ‘Protocols & publications’ tab (Fig. 5f). Here the users can find links to all publications linked to datasets currently contained within ImmPRes. In this same tab all the important details related the processing of the cells, the mass spectrometry and the analysis are also available for download as a PDF document.

## Code availability

ImmPRes has been developed in python and implemented via plotly (https://plotly.com/) dash (https://plotly.com/dash/). All of the proteomics data were processed using MaxQuant^37,38^ (https://www.maxquant.org/), with the search parameters included in each PRIDE submission. All copy numbers were calculated using R (https://www.r-project.org/) v 4.0.3, or with Perseus^39^ and the differential expression analysis performed with the Bioconductor package LIMMA^40^ v 3.50.3.

## Methods

### Mice

For proteomics experiments cell populations were isolated from C57BL/6 wild type mice, OT-I TCR transgenic ^41^ and P14 TCR transgenic mice^40^ and mice with T cell selective deletions of Myc ^42,43^or *Slc7a5*^29^ (full details are in the ‘Publications & protocols’ tab on immpres.co.uk). All mice for the Dundee based experiments were maintained in the Biological Resource Unit at the University of Dundee using procedures that were approved by the University Ethical Review Committee and under the authorization of the UK Home Office Animals (Scientific Procedures) Act 1986.

### Label free SP3 Proteomics sample preparation

Cell pellets composing of 1 – 8 million cells (depending on the number of fractions being analysed) were lysed in 400 μl lysis buffer at room temperature in 4% SDS, 50 mM TEAB pH 8.5, 10 mM TCEP under agitation (5 min, 1200 rpm on tube shaker), boiled (5 min, 500 rpm on tube shaker), then sonicated with a BioRuptor (30s on, 30s off x30 cycles). Protein concentration was determined using EZQ protein quantitation kit (Invitrogen) as per manufacturer instructions. Lysates were alkylated with 20 mM iodoacetamide for 1 hr at room temperature in the dark, before protein clean up by SP3^14^ procedure. Briefly, 200 µg of 1:1 mixed Hydrophobic and Hydrophilic Sera-Mag SpeedBead Carboxylate-Modified Magnetic Particles were added per protein sample then acidified to ∼pH 2.0 by addition 10:1 Acetonitrile: Formic Acid. Beads were immobilised on a magnetic rack and proteins washed with 2 × 70% ethanol and 1 × 100% acetonitrile. Rinsed beads were reconstituted in 0.1% SDS 50 mM TEAB pH 8.5, 1 mM CaCl2 and digested overnight with LysC followed by overnight digestion with Trypsin, each at a 1:50 enzyme to protein ratio. Peptide clean up was performed as per SP3 procedure^14^. Briefly, protein-bead mixtures were resuspended and 100% acetonitrile added for 10 min (for the last 2 min of this beads were immobilised on a magnetic rack). Acetonitrile and digest buffer were removed, peptides were washed with acetonitrile and eluted in 2% DMSO. Peptide concentration was quantified using CBQCA protein quantitation kit (Invitrogen) as per manufacturer protocol. Formic acid was added to 5% final concentration.

Samples were fractionated using high pH reverse phase liquid chromatography. Samples were loaded onto a 2.1 mm x 150 mm XBridge Peptide BEH C18 column with 3.5 μm particles (Waters). Using a Dionex Ultimate3000 system, the samples were separated using a 25 min multistep gradient of solvents A (10 mM formate at pH 9 in 2% acetonitrile) and B (10 mM ammonium formate pH 9 in 80% acetonitrile), at a flow rate of 0.3 mL/min. Peptides were separated into 16-32 fractions (depending on the experiment) which were consolidated into 8-16 fractions (depending on the experiment). Fractionated peptides were dried in vacuo then dissolved in 5% Formic Acid for analysis by LC-ES-MS/MS.

### Label free urea sample preparation

Cell pellets composing of 1 – 5 million cells (depending on the number of fractions being analysed) were lysed in 400 μl lysis buffer in urea buffer (8 M urea, 50 mM Tris pH 8.0, 1 mM TCEP) with protease (cOmplete mini EDTA free, Roche) and with phosphatase inhibitors (PhosStop, Roche) at room temperature for half an hour.

Samples were sonicated, and protein concentration was determined using a BCA assay according to the manufacturer’s instructions. To reduce samples, 10 mM DTT was added for 30 min at room temperature. After reduction, 50 mM iodoacetimide was added to each sample and incubated for 45 min at room temperature in the dark. Samples were then diluted to 4 M urea with Tris buffer (100 mM Tris pH 8.0, 1 mM CaCl2). LysC (Wako), reconstituted in Tris buffer, was then added to each sample at a ratio of 50:1 protein:LysC and the samples were incubated with LysC overnight at 30°C. Samples were then transferred to low-bind 15-ml falcon tubes (Eppendorf) and further diluted with Tris buffer to 0.8 M urea. Trypsin (Promega), reconstituted with Tris buffer, was then added to the samples at a ratio of 50:1 protein:trypsin and the samples were incubated for 8 hours at 30°C. After digestion, the samples were desalted with C18 SepPack cartridges (Waters) and dried down in a SpeedVac (Genevac). Dried peptide samples were resuspended in 10 mM sodium borate 20% (v/v) acetonitrile (pH 9.3) and fractionated into 16 fractions by strong anion exchange (SAX) chromatography. Peptide samples for proteomic analysis were fractionated with an Ultimate 3000 HPLC equipped with an AS24 strong anion exchange (SAX) For the separation, the buffers used were 10 mM sodium borate (pH 9.3) (SAX buffer A) and 10 mM sodium borate (pH 9.3), 500 mM NaCl (SAX buffer B). Peptide samples were resuspended in 210 µl of 10 mM sodium borate 20% (v/v) acetonitrile (pH 9.3) and injected onto the SAX column and separated using an exponential elution gradient starting with Buffer A. In total, 16 peptide fractions were collected and desalted with Sep-pack C18 96 well desalting plates (Waters). Desalted peptides were dried down with a SpeedVac (Genevac).

### Label Free Liquid chromatography tandem mass spectrometry analysis

For each fraction, 1 μg was injected onto a nanoscale C18 reverse-phase chromatography system (UltiMate 3000 RSLC nano, Thermo Scientific) then electrosprayed into an Orbitrap mass spectrometer (LTQ Orbitrap Velos Pro; Thermo Scientific). For chromatography buffers were as follows: HPLC buffer A (0.1% formic acid), HPLC buffer B (80% acetonitrile and 0.08% formic acid) and HPLC buffer C (0.1% formic acid). Peptides were loaded onto an Acclaim PepMap100 nanoViper C18 trap column (100 µm inner diameter, 2 cm; Thermo Scientific) in HPLC buffer C with a constant flow of 10 µl/min. After trap enrichment, peptides were eluted onto an EASY-Spray PepMap RSLC nanoViper, C18, 2 µm, 100 Å column (75 µm, 50 cm; Thermo Scientific) using the buffer gradient: 2% B (0 to 6 min), 2% to 35% B (6 to 130 min), 35% to 98% B (130 to 132 min), 98% B (132 to 152 min), 98% to 2% B (152 to 153 min), and equilibrated in 2% B (153 to 170 min) at a flow rate of 0.3 µl/min. The eluting peptide solution was automatically electrosprayed using an EASY-Spray nanoelectrospray ion source at 50° and a source voltage of 1.9 kV (Thermo Scientific) into the Orbitrap mass spectrometer (LTQ Orbitrap Velos Pro; Thermo Scientific). The mass spectrometer was operated in positive ion mode. Full-scan MS survey spectra (mass/charge ratio, 335 to 1800) in profile mode were acquired in the Orbitrap with a resolution of 60,000. Data were collected using data-dependent acquisition: the 15 most intense peptide ions from the preview scan in the Orbitrap were fragmented by collision-induced dissociation (normalized collision energy, 35%; activation Q, 0.250; activation time, 10 ms) in the LTQ after the accumulation of 5000 ions. Precursor ion charge state screening was enabled, and all unassigned charge states as well as singly charged species were rejected. The lock mass option was enabled for survey scans to improve mass accuracy. (Using Lock Mass of 445.120024).

### TMT Proteomics sample preparation

Cell pellets composing of 1 – 5 million cells (depending on the population that was analysed) were lysed in 400 μl lysis buffer (4% sodium dodecyl sulfate, 50 mM TCEP (pH 8.5) and 10 mM tris(2-carboxyethyl)phosphine hydrochloride). Lysates were boiled and sonicated with a BioRuptor (30 cycles: 30 s on and 30 s off) before alkylation with 20 mM iodoacetamide for 1 h at 22 °C in the dark. The lysates were subjected to the SP3^14^ procedure for protein clean-up^14^ before elution into digest buffer (0.1% sodium dodecyl sulfate, 50 mM TEAB (pH 8.5) and 1 mM CaCl_2_) and digested with LysC and Trypsin, each at a 1:50 (enzyme:protein) ratio. TMT labeling and peptide clean-up were performed according to the SP3 protocol.

The TMT samples were fractionated using off-line high-pH reverse-phase chromatography: samples were loaded onto a 4.6 mm × 250 mm XbridgeTM BEH130 C18 column with 3.5 μm particles (Waters). Using a Dionex BioRS system, the samples were separated using a 25-min multistep gradient of solvents A (10 mM formate at pH 9 in 2% acetonitrile) and B (10 mM ammonium formate at pH 9 in 80% acetonitrile), at a flow rate of 1 ml min^-1^. Peptides were separated into 48 fractions, which were consolidated into 24 fractions. The fractions were subsequently dried, and the peptides were dissolved in 5% formic acid and analyzed by liquid chromatography–mass spectrometry.

### TMT Liquid chromatography electrospray–tandem mass spectrometry analysis

For each fraction, 1 μg was analysed using an Orbitrap Fusion Tribrid mass spectrometer (Thermo Fisher Scientific) equipped with a Dionex ultra-high-pressure liquid chromatography system (RSLCnano). Reversed-phase liquid chromatography was performed using a Dionex RSLCnano high-performance liquid chromatography system (Thermo Fisher Scientific). Peptides were injected onto a 75 μm × 2 cm PepMap-C18 pre-column and resolved on a 75 μm × 50 cm RP C18 EASY-Spray temperature-controlled integrated column-emitter (Thermo Fisher Scientific) using a 4-h multistep gradient from 5% B to 35% B with a constant flow of 200 nl min^-1^. The mobile phases were: 2% acetonitrile incorporating 0.1% formic acid (solvent A) and 80% acetonitrile incorporating 0.1% formic acid (solvent B). The spray was initiated by applying 2.5 kV to the EASY-Spray emitter, and the data were acquired under the control of Xcalibur software in a data-dependent mode using the top speed and 4 s duration per cycle. The survey scan was acquired in the Orbitrap covering the *m/z* range from 400–1,400 Thomson units (Th), with a mass resolution of 120,000 and an automatic gain control (AGC) target of 2.0 × 10^5^ ions. The most intense ions were selected for fragmentation using collision-induced dissociation in the ion trap with 30% collision-induced dissociation energy and an isolation window of 1.6 Th. The AGC target was set to 1.0 × 10^4^, with a maximum injection time of 70 ms and a dynamic exclusion of 80 s. During the MS3 analysis for more accurate TMT quantifications, ten fragment ions were co-isolated using synchronous precursor selection, a window of 2 Th and further fragmented using a higher-energy collisional dissociation energy of 55%. The fragments were then analyzed in the Orbitrap with a resolution of 60,000. The AGC target was set to 1.0 × 10^5^ and the maximum injection time was set to 300 ms.

### Processing and analysis of proteomics data

All the datasets were processed, searched and quantified with MaxQuant^37,38^. The search parameters for the individual datasets are included in each pride submission, however in all datasets the false discovery rate was set to 1% for positive identification at the protein and at the peptide-spectrum match level.

### Copy number calculations

The estimated protein copy numbers were calculated using the proteomic ruler^17^. The TMT derived estimated copy numbers required additional steps which involved assigned the MS1 intensity based on the ratios of the MS3 reporter ion intensity as previously described^18^.

### Differential expression analysis

All the differential expression analysis on ImmPRes were calculated in R based on the estimated protein copy numbers and using the Bioconductor package limma^40^.

### Figure generation

The bar plots in this paper were generated in R. The schematics were all generated using biorender.com and Sankey diagram was generated using SankeyMATIC (https://sankeymatic.com/)

## Acknowledgements

We would like to thank Colin Watts for contributing the eosinophil population within the ‘Hematopoietic cell proteomes’ dataset, all the members of the Lamond and Cantrell labs for their feedback and discussions relating to the design and testing of ImmPRes and the Fingerprint Proteomics Facility at the University of Dundee, for their support in processing the label-free DDA samples. This work was funded by a Wellcome Trust strategic award (105024/Z/14/Z) to AIL and DAC, Wellcome Trust grants (097418/Z/11/Z & 205023/Z/16/Z) to DAC, Wellcome Trust grant (073980/Z/03/Z) to AIL, a Wellcome Trust Equipment Award (202950/Z/16/Z) and a UK Research Partnership Infrastructure Fund award to the Centre for Translational and Interdisciplinary Research. JMM was supported by funding from the European Union’s Horizon 2020 research and innovation programme under the Marie Skłodowska-Curie grant agreement No 705984 awarded to JMM and DAC.

## Notes

### Competing Interest Statement

The authors have declared no competing interest.

